# Distributed dynamic coding for spatial working memory in hippocampal-prefrontal networks

**DOI:** 10.1101/630673

**Authors:** AE Hernan, JM Mahoney, W Curry, S Mawe, RC Scott

## Abstract

Spatial working memory (SWM) is a central cognitive process during which the hippocampus and prefrontal cortex (PFC) encode and maintain spatial information for subsequent decision-making. This occurs in the context of ongoing computations relating to spatial position, recall of long-term memory, attention, amongst many others. To establish how intermittently presented information is integrated with ongoing computations we recorded single units, in both hippocampus and PFC, in control rats and those with a brain malformation during performance of a SWM task. Neurons that encode intermittent task parameters are also well-modulated in time and incorporated into a functional network across regions. Our results implicate a model in which ongoing oscillatory coordination among neurons in the hippocampal-PFC network defines a functional network that is poised to receive sensory inputs that are then integrated and multiplexed as working memory. These dynamics are systematically altered in disease and may provide potential targets for stimulation-based therapies.

## Introduction

Spatial working memory (SWM) is the cognitive process by which goal-related information from the external world is encoded, maintained and integrated by the brain so that it can be accessed by neural circuits that plan ahead and execute goal-oriented behavior^1^. Current models posit that accurate SWM requires functional coordination of distributed networks throughout the brain^2,3^. In particular, the hippocampus encodes the spatial position of the relevant stimulus and sends the information to the prefrontal cortex (PFC) where it is dynamically maintained and ultimately used for decision-making^4–9^. Importantly, this has to be performed in the context of multiple other ongoing computations involving motivation, recall of long-term memory, attention and possible motor strategies amongst many others. Therefore, SWM requires a *distributed dynamic code*, in which a stimulus elicits a dynamical pattern of neuronal firing that is robustly associated with the stimulus’ identity, is maintained throughout goal-oriented computation^10^ and that is integrated with ongoing computations. Disruptions to the neural circuits that generate and maintain these distributed dynamical codes could represent a system-level mechanism underpinning the SWM deficits frequently observed, for example, in epilepsy and schizophrenia^11–13^.

The hippocampal-prefrontal network is a continuously operating system that has this capacity to encode and maintain specific information during an SWM task in concert with ongoing computations.^7,14,15^. The neural circuits supporting a SWM task that has already been learned must have multiple, latent dynamical patterns corresponding to distinct task computations that can each be elicited by appropriate stimuli, and that are integrated with all other functions. Thus, the functional architecture of the hippocampal-prefrontal network, i.e. the coordinated firing of cells in time with respect to each other encapsulates all computation during the task. The relationships between the co-firing behavior among cells throughout a SWM session (consistent with ongoing computation of the multiple unmeasured aspects of the task) and transient responses related to stimulus-perception and decision-making remain uncertain. At the level of single unit activity in the hippocampus, there are well-established links between oscillatory firing on one hand and rate coding of external outputs on the other hand. For example, hippocampal pyramidal cells are strongly theta modulated during navigation, and this is strongly correlated with the spatial tuning of firing^16–18^. In SWM, *in silico* evidence suggests that ongoing oscillatory modulation is critical for building appropriate rate coding dynamics, especially when crucial decision-points are unreliably timed^19^. Beyond oscillatory firing at the single neuron level within the PFC, it is clear that the hippocampal-prefrontal network also has tightly regulated coordination between structures at the level of local field potentials, including coherence in multiple frequency bands^6,20–22^. We hypothesized, therefore, that the neurons in the hippocampal-prefrontal network that encode SWM also have tightly regulated firing and co-firing relationships, defining a distributed dynamic code for the SWM task, and that SWM deficits in disease correspond to alterations in the topology of these functional networks.

We investigated the functional behavior of single units in the PFC and CA1 of the hippocampus during a delayed non-match-to-sample (DNMS) task. We studied normal rats to define the relationships in physiology and rats that were exposed to methylazoxymethanol (MAM) *in-utero* to identify differences in a structurally abnormal brain. MAM administration generates a clinically-relevant model of brain malformation that persists into adulthood^23,24^. This structural outcome is consistent with human malformations of cortical development caused by many genetic and environmental insults, and which are known to negatively affect working memory. We used a rigorous generalized linear modeling (GLM) approach to systematically assess the cognitive, oscillatory, and functional network properties of simultaneously recorded neural ensembles. We show that the neurons that encode task parameters at decision points during the task are the same neurons whose firing is well-modulated in time throughout the duration of an experimental session, consistent with *in-silico* models. These neurons are substantially more likely to be functionally connected to other neurons both within and between brain regions. Moreover, these functional networks are strongly enriched for neurons in both regions that multiplex information about both the sample lever position as well as the match phase choice, suggesting that the acquisition, maintenance, and decision-making patterns of activity are all supported by a tightly integrated functional network. MAM animals have poorer background firing modulation in the hippocampus, fewer neurons that encode task parameters, and fewer functionally connected pairs of neurons. These alterations directly predict task performance. Thus, the functional network structure is a fundamental property of hippocampal-PFC networks that behaves like a scaffold upon which the task-related dynamics holding memories are built. These background dynamics are systematically altered in disease and may provide potential therapeutic targets for stimulation-based therapies in the future.

## Results

To evaluate the relationships between overall neural dynamics and task related dynamics measured during a working memory task, and to understand how these dynamics are altered in developmentally abnormal networks, we simultaneously recorded single units in the PFC and the CA1 region of the dorsal hippocampus of control and an embryonic-day-17 methylazoxymethanol (MAM) model during performance of a DNMS task (Fig 1). In six control and seven MAM rats, a total of 739 cells were simultaneously recorded from both brain regions during 5-10 sessions of the DNMS task. Task delay lengths were increased each day in an overlapping sliding window and ranged from 0s to 30s during the course of the experiment.

**Figure 1.**
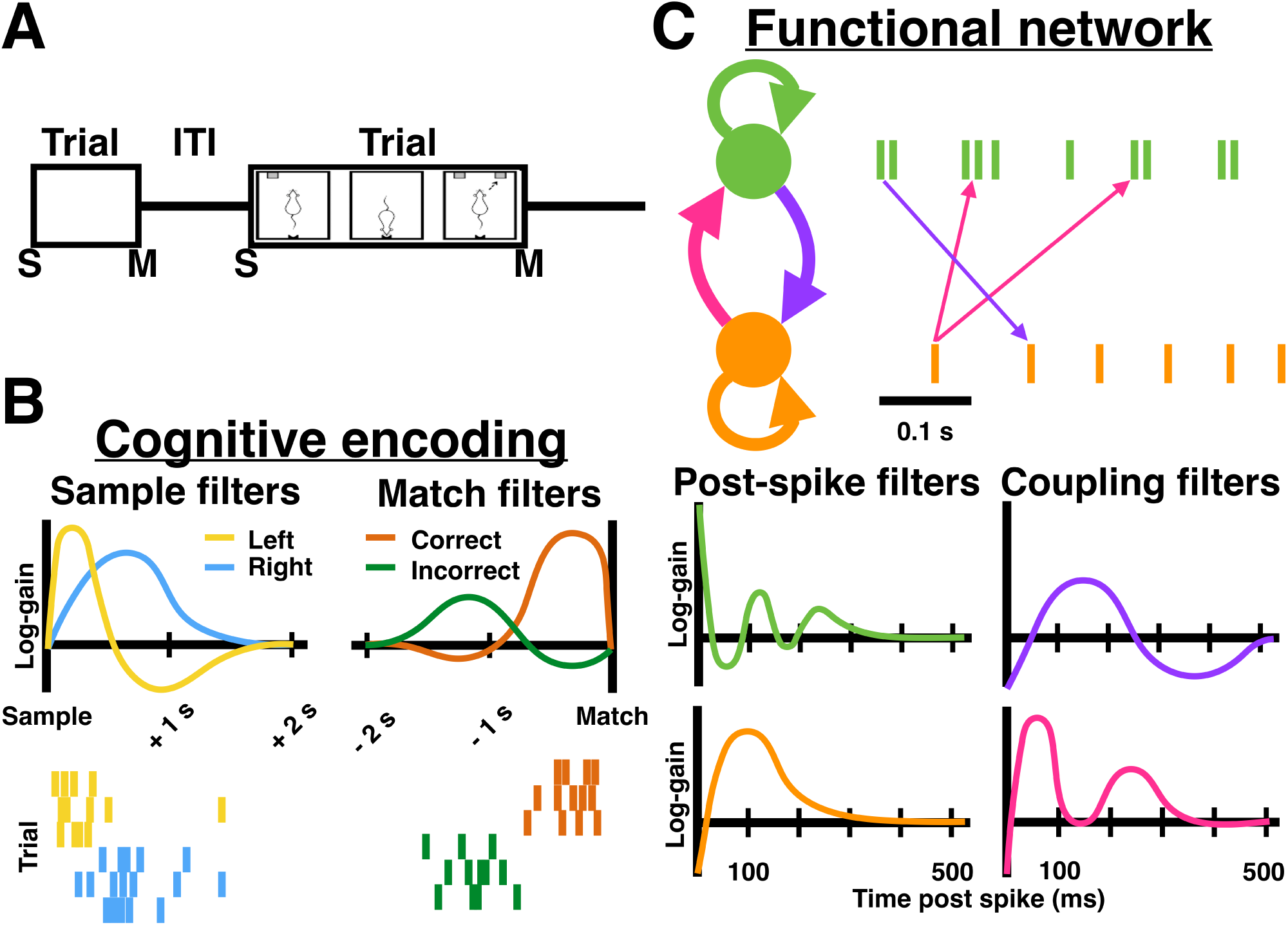
Generalized linear modeling (GLM) approach to cognitive encoding and functional networks. A) We trained animals to perform a delayed non-match to sample (DNMS) task. At the beginning of each trial, a single lever, either left or right, is presented to the rat (sample phase). After a variable delay, the rat is prompted to break a laser beam with its nose at the back of the chamber (nosepoke). Immediately after the nosepoke, both levers are presented and the rat is rewarded if it presses the opposite lever to the sample (match phase). B) To rigorously measure cognitive encoding at the sample and match phases, we modeled neuronal firing around each lever press using a GLM. For approximately 2 s after the sample press and before the match press, the GLM modeled the probability of firing as a smooth modulation of the background firing rate (*sample* and *match filters*). If the filter value is positive (resp. negative), then the neuron has a much higher (resp. lower) probability of firing relative to its baseline rate. In these toy examples, the neuron fires sooner after a left lever press than a right press, and closer to the match press in correct trials than incorrect trials. The sample and match filters are, therefore, effectively smoothed peri-event histograms. C) To rigorously measure functional connectivity among neurons throughout the session, we used GLMs that captured auto-correlations and cross-correlations of neural firing. Auto-correlation is captured by a *post-spike filter* (PSF), which modifies the future firing probability after a spike. For example, the spike train of the green cell has a bursting phenotype with prominent theta oscillation, which is modeled by the PSF as a pronounced up-regulation immediately post spike followed by another up-regulation 0.1 s later. In contrast, the orange cell has a regular firing phenotype with a refractory period followed by an up-regulation at 0.1 s and a decay to baseline. Cross-correlations are captured by *coupling filters* (CPFs), which modify the future firing probability of a neuron after a spike of another neuron. For example, a spike of the green neuron predicts an immediate decrease in probability of the orange neuron firing, followed by an increase 0.05-0.25 s post spike, and a subsequent decrease from 0.25-5 s post spike (purple arrows and filter). Conversely, a spike of the orange neuron predicts an increased probability of the green neuron firing 0.05 s and 0.15 s later (pink arrows and filter).

### MAM rats have behavioral deficits that can be partially ameliorated by training

We assessed the behavior of adult MAM rats compared to saline-exposed controls during DNMS. Previous work by our group has shown that MAM rats can acquire spatial and non-spatial tasks with training^25^, so we utilized a training protocol that ensured that MAM rats were able to perform the working memory component of the DNMS task during recording sessions. In line with our previous observations, MAM rats take significantly longer to reach criterion compared to controls (Fig. 2a, p<0.001), but once the task was acquired MAM rats were able to accurately perform over delays ranging from 0s to 30s (Fig. 2b). As expected, in both groups of animals there was a clear reduction in performance with increasing delays (p<0.001). Despite performing as accurately as their control counterparts, MAM animals took longer to complete the task. While output delay lengths from the behavioral software were the same by session between the groups (Fig. 2c), MAM rats took on average 1.7s (11.53s, CI: 10.15-13.09s) longer than control rats (9.85s, CI: 8.82-11.0s) to complete each correct, but not incorrect, trial (p=0.007; Fig. 2d). This is also reflected as an increase in total time to complete each session (Fig. 2e).

**Figure 2.**
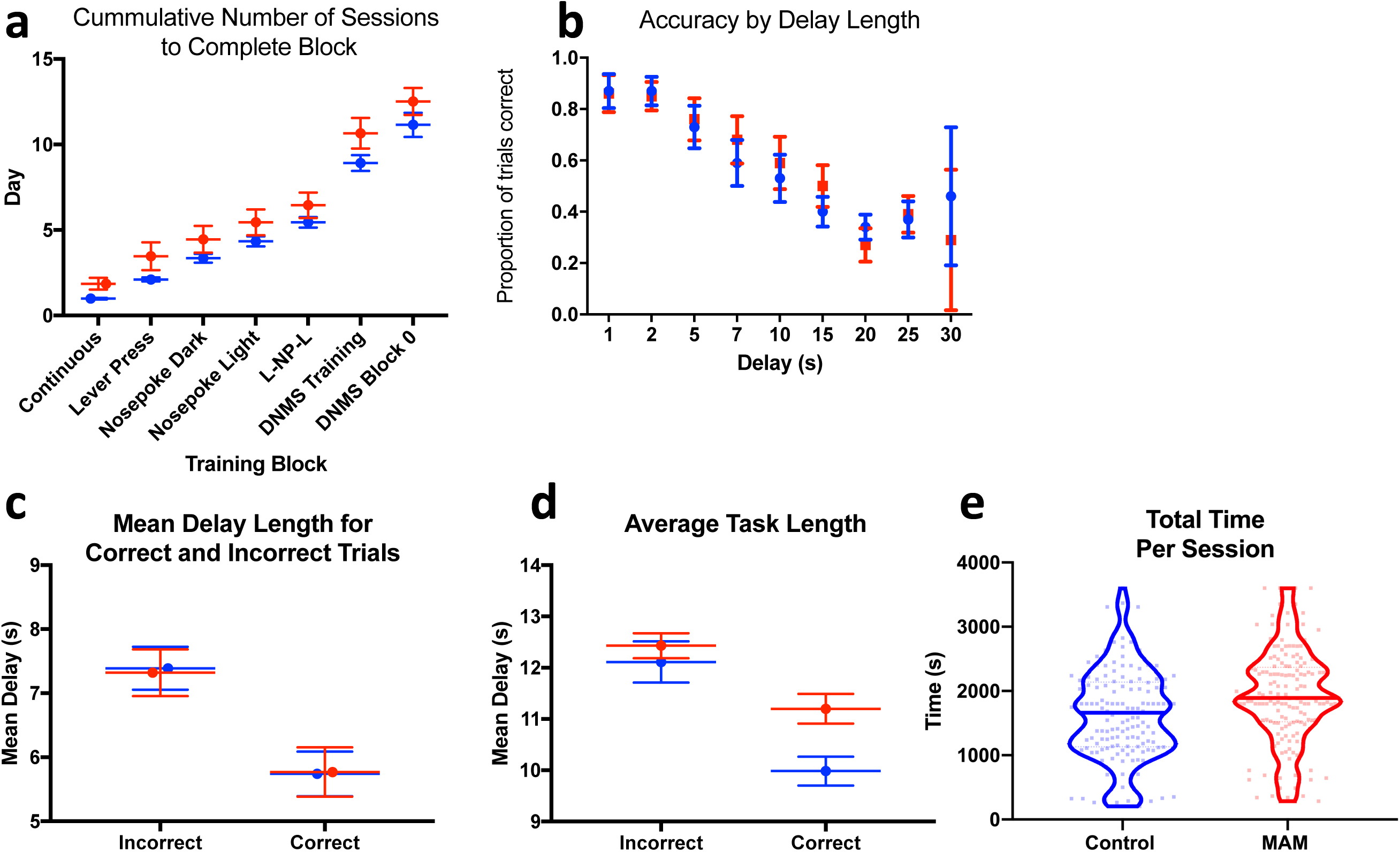
MAM animals perform the SWM task as well as controls but require more time to make a decision. MAM animals (shown in red) took significantly longer than controls (shown in blue) to acquire the DNMS task (a). Once they were fully trained, they were able to perform as accurately as controls at delays ranging from 1-30s (b). However, despite seeing identical programmed delay lengths (c), MAM animals took longer to make correct, but not incorrect, choices than their control counterparts (d). This manifests as a slight but significant increase in overall session length in the MAM animals (e).

Thus, overtraining MAM rats recovers some working memory function, but does not completely restore performance to normal. Because of these differences, we hypothesized that MAM animals had abnormal neural firing relative to controls, corresponding to altered cognitive processing during the task. We therefore studied single unit activities at decision points (cognitive encoding), throughout the entire session on a cell-by-cell basis (oscillatory timing), and at the level of interactions among cells within and between brain structures (functional networks).

### Rate modulation around the sample and match press indicate differences in encoding of the levers across groups

To quantify dynamical patterns of neural firing with limited *a priori* assumptions about firing modulation, we used a GLM approach. GLMs are a statistically principled way to model spiking activity using flexible parametric representations that capture heterogeneity across neurons^26,27^ and allow for rigorous statistical hypothesis testing ^28^.

Our initial analyses evaluated task-related neural dynamics during the performance of the behavioral components of the DNMS task, i.e. during the two seconds after the sample lever press and the two seconds prior to the match lever press. The GLM encodes the modulation of firing rate around these events using an *event filter*, which is a time varying curve that tracks the *gain* of firing rate relative to the baseline (Poisson) firing rate (Fig 1b, Fig 3). In this way, the event filter is a parametric model of the peri-event time histogram (PETH) transformed to log-space to capture fold changes in the rate of firing (Fig 3; see also Methods). Event filters confirm heterogeneity of firing shown in previous experiments (Supplemental Fig.1). To establish whether a neuron specifically discriminated between left and right lever presses, we compared GLMs with distinct event filters for left and right lever presses (left-right model) to another that had only one filter for all lever presses (null model). The null model allows rate modulation at task performance times, but does not discriminate between levers, and is therefore a null model for whether the neuron encodes the task. We then computed the log-likelihood ratio (LLR) between these models and used a rigorous statistical cutoff to establish whether the left-right model better accounts for firing activity relative to the null model. Neurons that had a significant LLR at the sample phase were defined as *encoding the sample*. We find that fewer neurons in both the hippocampus and the PFC of MAM animals compared to controls encoded the sample (Fig 3a,b; 54% in controls vs 48% in MAM in the hippocampus, p=0.006; 59% in controls vs 44% in MAM in the PFC, p<0.001)

**Figure 3.**
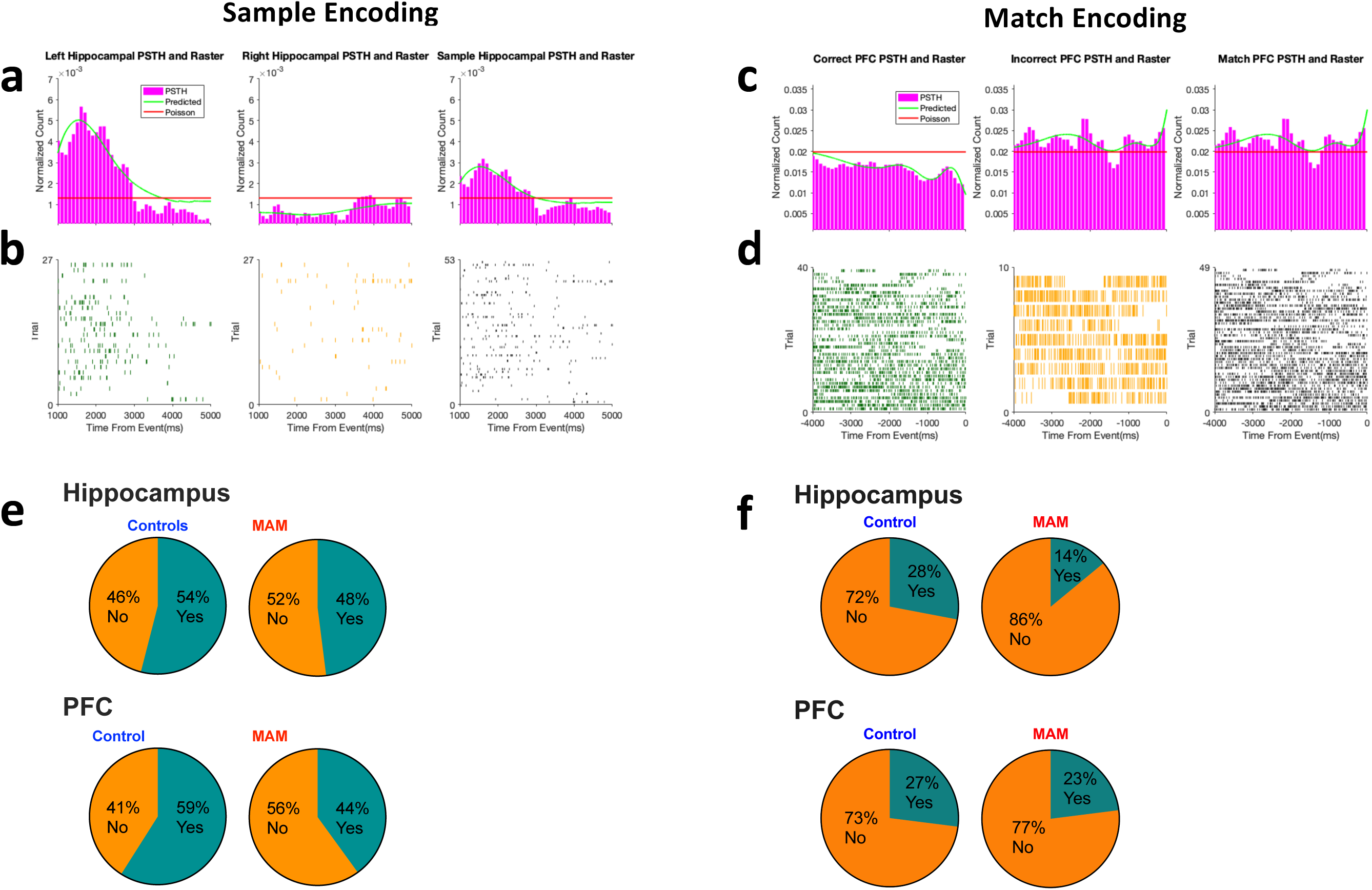
Neurons in both the hippocampus and the PFC encode sample and match choices. Data on the left (a, b, e) are from the sample phase, right panels (c, d, f) are data from the match phase. Representative binned histograms in pink from the left sample press and the right sample press with the sample event filter fit overlaid in green (a). Raster plots of a single neuron in the hippocampus whose firing discriminates left lever press from right lever press trials (b). Likewise, representative binned histograms in pink from the correct and the incorrect choice points at the match phase with the match event filter fit overlaid in green (c). Raster plots of a single neuron in the PFC whose firing discriminates correct from incorrect trials (d). Both of these neurons have significant LLRs for sample and match presses, respectively, indicating good encoding of the task parameter. Group average pie charts show proportions of neurons that significantly encode the sample lever position in the hippocampus (top pie charts, e) and PFC (bottom pie charts, e). Group average pie charts show proportions of neurons that significantly encode the match phase outcome in the hippocampus (top pie charts, f) and PFC (bottom pie charts, f).

Similarly, using event filters for dynamics from 2s prior to the match phase, we computed LLRs to determine whether neurons from each brain region in MAM and controls significantly distinguished between correct and incorrect levers. Rate coding with respect to the match phase lever shows a similar pattern to the data from the sample phase. Fewer hippocampal neurons in MAM rats (28% vs 14%; p<0.001) distinguish between a correct and an incorrect lever press (Fig. 3e,f); however, a similar proportion of neurons (27% vs 23%, p=0.25) from the PFC encode correct vs incorrect trials (Fig. 3e,f). To characterize this relationship further, we categorized neurons by whether they significantly encoded the sample, the match phase choice, neither, or both. Any given cell in the hippocampus of controls is 1.7 times (95% CI;1.6-1.8) more likely to encode both the sample and the match phase than in MAM animals (p<0.001). Any given cell in the PFC of controls is 2.0 times (95% CI;1.4-2.9) more likely to code both the sample and the match than in MAM animals (p<0.001). This demonstrates that the subset of neurons incorporated into the task-encoding ensemble is smaller, suggesting the network as a whole is of lower fidelity in MAM animals compared to controls.

There is a significant relationship between the proportion of cells that have significant LLRs at either the sample or the match phase and overall accuracy in performance, after accounting for group, mean trial length and interactions between LLRs and trial length. The model accounts for 54% of the variance (P<0.001, Supplemental Fig. 2a) confirming that task related neural dynamics in the hippocampus and PFC are critically related to the performance of the task.

In addition to fewer numbers of neurons encoding the match phase in MAM animals, we also examined the temporal patterning of firing underlying correct performance on a cell-by-cell basis. To do this, we subtracted the filters of incorrect presses from the filters of correct presses to examine the difference in firing modulation (Fig. 4). This establishes a striking pattern of differential firing between MAM and control animals, both in the hippocampus (Fig. 4A) and the PFC (Fig. 4B). In the hippocampus, there are approximately equal numbers of neurons whose firing is up- or down-regulated in correct relative to incorrect trials. This relative up- or down-regulation of firing occurs around 2s in advance of the match choice. In MAM animals, the population of neurons whose firing was down-regulated in correct trials compared to incorrect trials is practically eliminated. In addition, the up-regulation occurred immediately (<1s) before the match choice. In the PFC, this loss of down-regulated neurons is less pronounced and the up-regulation of firing occurs both at 2s and <1s immediately preceding the match in MAM animals, but is only seen at 2 seconds before the match in controls (Fig. 4). This suggests group differences in the ensemble-level dynamics related to solving the task. We next investigated the ongoing oscillatory and functional network structure among neurons to identify the network properties that determined whether a cell encoded task parameters, and how these properties differed between groups.

**Figure 4.**
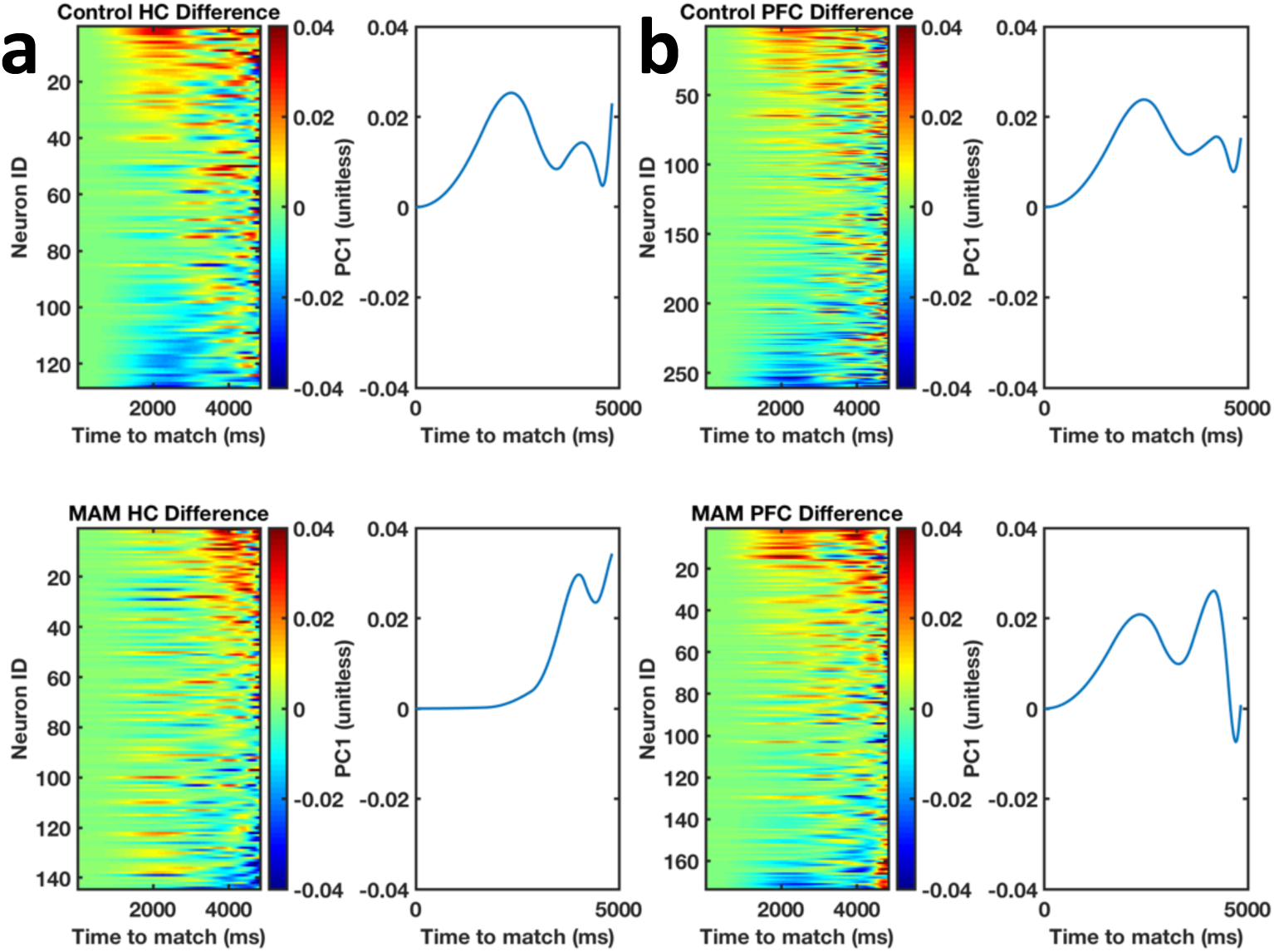
Neuronal firing in advance of a match press is different between the groups. Heatmaps of the incorrect minus the correct event filters 3s before the match decision point are shown. Heatmaps were sorted by pattern of firing as determined by a principal component analysis (PCA), with the first principal component (PC 1) shown to the right of each heatmap. Control hippocampal neurons (a) and PFC neurons (b) show two patterns of firing: a population that upregulates its firing approximately 2 s before the decision point, and a smaller population that downregulate their firing 2 s before a decision point. In MAM animals, hippocampal neurons (a) and PFC neurons (b) have shifted their firing to approximately 500 ms before the choice point, and the population of downregulated neurons has largely been lost.

### Temporal modulation of ongoing fine spike timing differs between groups and predicts rate encoding of task parameters

To evaluate the short-timescale modulation (in the order of milliseconds) of fine spike timing (FST), we modeled spike trains using a GLM that incorporated the spiking history of the neuron to predict future firing. In this model, FST properties are captured by *post-spike filters* (PSF), which, similar to the event filters, encodes the gain in firing rate as a function of past spiking^26^ (Fig. 1C). The PSF allows the GLM to accurately model the auto-correlation of firing over the entire session during the DNMS task, including periods when the animal is actively engaged in the task as well as inter-trial periods. To quantify differences in FST between the groups in an unbiased manner, we performed a principal component analysis (PCA) on the PSFs and used the first principal component (PC 1) score to evaluate the fidelity of short timescale modulation of hippocampal and prefrontal neuronal populations (Fig. 5).

**Figure 5.**
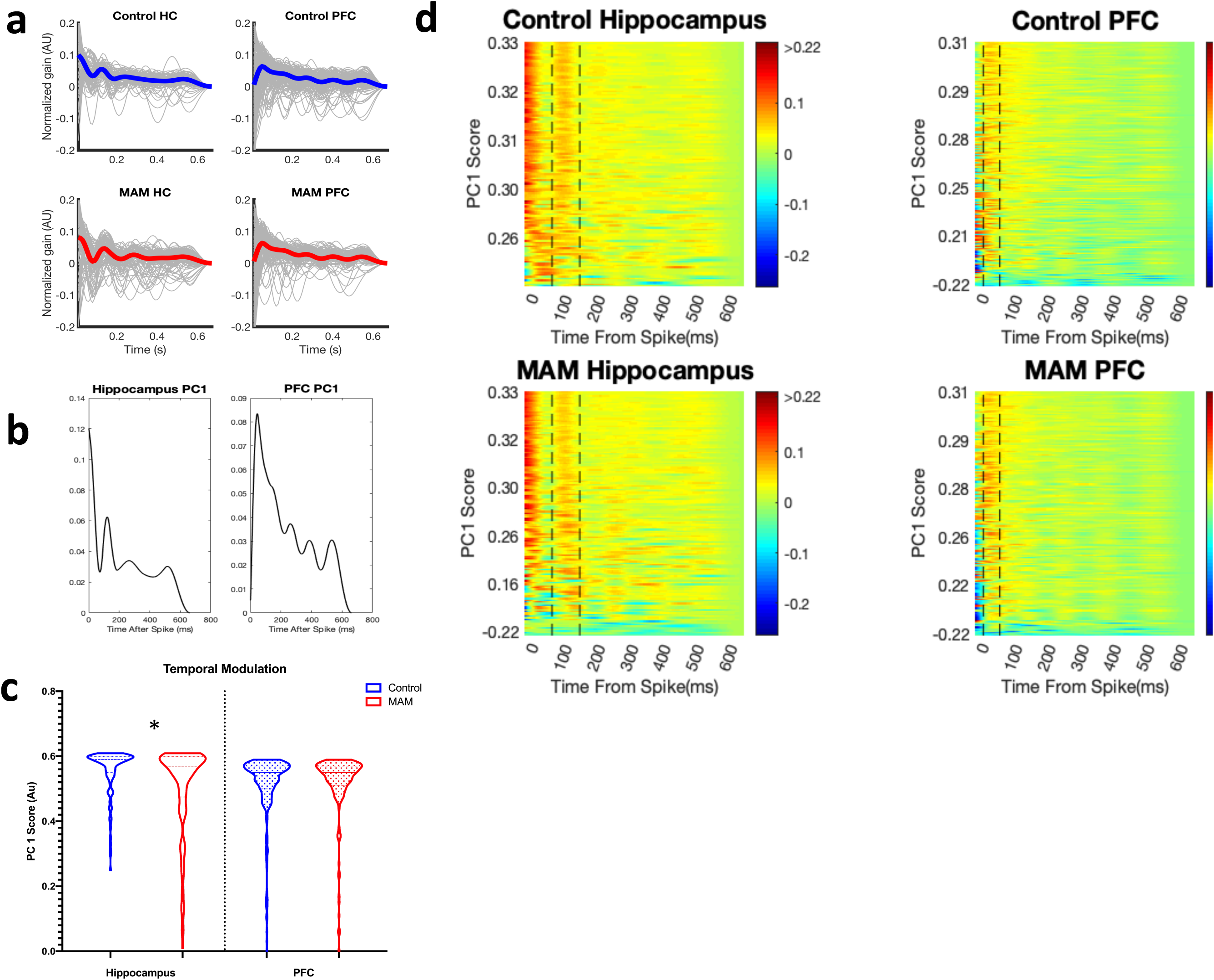
Fine spike timing temporal modulation of hippocampal, but not PFC neurons is poorer in MAM animals. Postspike filter averages (dark lines) are overlaid against all individual neurons (gray lines) for control hippocampus (top left in blue, **a**), control PFC (top right in blue, **a**) and MAM hippocampus (bottom left in red, **a**), MAM PFC (bottom right in red, **a**). This overall pattern is recapitulated in the PC 1 of hippocampus (right, **b**) and PFC (left, **b**). PCA allows for unbiased quantification of the amount of temporal modulation between groups (**c**), with hippocampal neurons from MAM animals (red) showing lower median PC 1 scores than controls (blue) in the hippocampus but not the PFC. Heat maps of the entire population reveal a strong theta modulation in control (top left, **d**) and MAM hippocampus (bottom left, **d**) as seen as warmer colors corresponding to upregulation of firing probability between dotted lines representing 6 and 12 Hz. The PFC from control neurons (top right, **d**) and MAM neurons (bottom right, **d**), a different pattern of fine spike timing temporal modulation than hippocampal neurons, with an initial upregulation shown as warmer colors between the dotted lines at 20-50ms, although no significant differences were seen in temporal modulation of PFC neurons between MAM and control.

In the hippocampus, PC 1 shows a strong upregulation of the probability of firing within the first 10 ms after a spike, with another upregulation 120 ms later, corresponding to 8.3 Hz modulation (theta modulation; Fig. 5). This captures typical firing dynamics of hippocampal neurons with burst firing and theta modulation^29^. MAM neurons in the hippocampus have lower average PC 1 scores (Fig. 5b, c; 0.56 ± 0.004 in control neurons vs 0.50 ± 0.011 in MAM neurons, p<0.001), indicating less precise theta modulation. In the PFC, the first principal component shows a strong immediate downregulation after an initial spike, with an upregulation at 20-50 ms after the initial spike, corresponding to a regular firing phenotype, typical of cortical neurons^30^ (Fig. 5d). No differences were noted between groups in PC 1 scores for PFC neurons. Thus, there are clear overall oscillatory abnormalities in the hippocampi of MAM animals.

It is not *a priori* obvious that the neurons encoding task parameters ought to also have specific oscillatory timing characteristics. However, there is *in-silico* evidence suggesting a relationship between timing dynamics and generation of appropriate rate dynamics. We therefore evaluated whether the post-spike filter PC1 score recorded from an individual neuron predicted whether or not *that* neuron was significantly encoding task parameters (i.e. significant sample or match LLRs). PC 1 scores were converted to *normal scores* by taking the inverse normal cumulative distribution function of their normalized ranks^31^. For each standard deviation increase in the normal scores, a hippocampal neuron was 1.3 times (95% CI; 1.1-1.6) more likely to discriminate between the right and left lever at the sample phase (p=0.01, Fig 6) and 1.3 times (95% CI; 1.06 −1.6) more likely to discriminate correct from incorrect trials (p=0.01). There were no significant relationships between the PC1 score in the PFC and significant encoding of task parameters. Taken together, these data indicate that overall oscillatory firing of hippocampal neurons throughout the session predicts whether an individual cell will encode intermittently presented task parameters (Supplemental Fig 2b).

**Figure 6.**
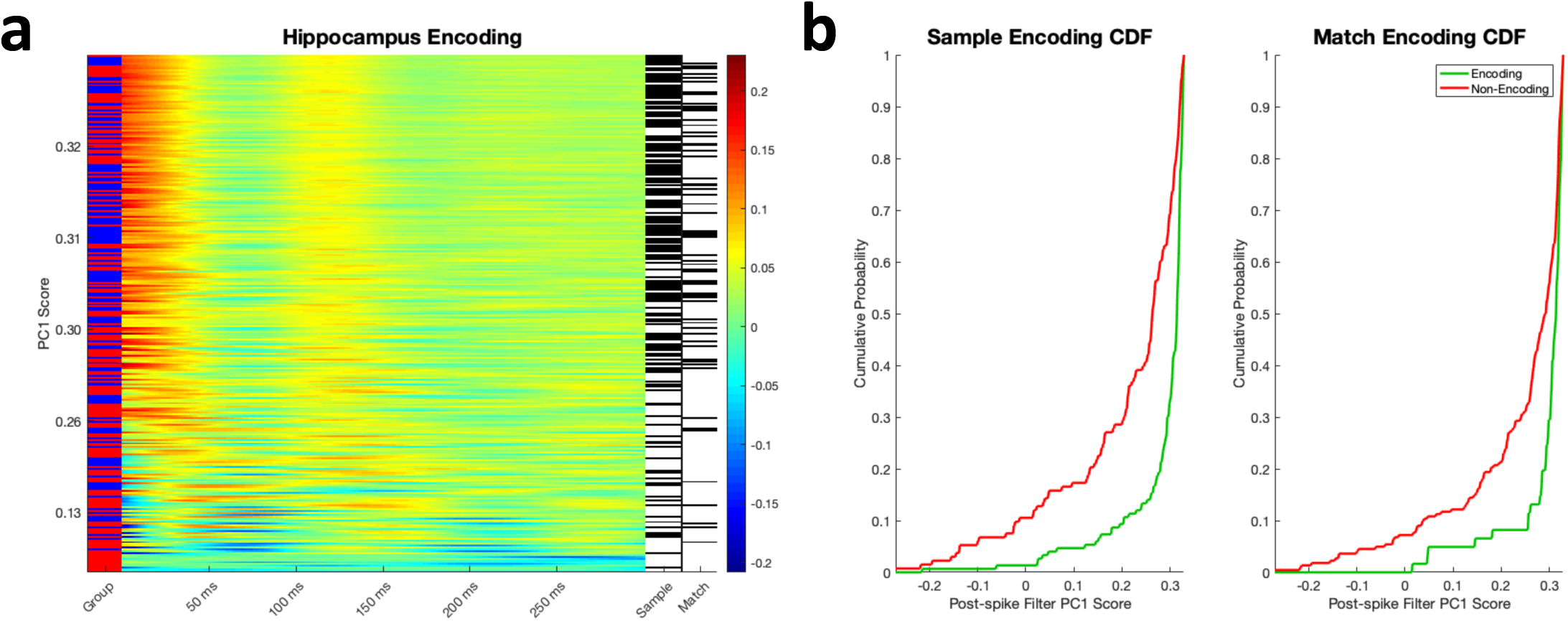
Hippocampal neurons that encode the sample and the match on the second timescale are the same neurons that are well-temporally modulated in the millisecond timescale. Heatmaps of the postspike filters are shown for all hippocampal neurons (a) and PFC neurons (b). Neurons from control animals are labeled with blue in the left-hand column MAM animals are labeled with red in the left-hand column of each heat map. Hippocampal neurons that significantly encode the sample and the match (a) are labeled in black on the right-hand column of the heatmap; neurons that do not encode are labeled in white. Cumulative distribution function graphs are shown in as a quantification of the observation made in the heatmaps that neurons that encode the sample in the hippocampus (green trace in the left graph, b) have higher PC 1 scores (e.g. better temporal modulation) than neurons that do not encode, shown in red. Likewise, hippocampal neurons that encode the match (green trace in the right graph, c) have higher PC 1 scores (e.g. better temporal modulation) than neurons that do not encode, shown in red.

### Fidelity of the hippocampal-PFC functional network predicts rate coding and performance

The above analyses concentrated on the behavior of single cells. However, these cells are likely to be coordinated within functional networks, the character of which could be important for SWM performance. To identify functional networks within the measured ensembles, we jointly modeled spike trains using a GLM that included both PSFs and *coupling filters* (CPFs), which encode the gain in firing rate as a function of past spiking of *other* neurons in the ensemble. A CPF corresponds to a functional link between neurons, by which a spike of the source neuron predicts a corresponding modulation of the firing probability of the target neuron (Fig. 1c). Importantly, CPFs are fitted simultaneously, so that each CPF models the specific firing modulation of the target neuron relative to the source neuron after accounting for the firing of all other neurons^26^. Given the large number of parameters in the coupled models, we used Granger causality (*i.e.*, an LLR test) to determine whether pairs of cells were significantly comodulated across the entire session^28^.

We defined a functional network for the ensemble by adding an edge for every significant CPF among the cells. Edges were identified in 55% of all possible hippocampal to hippocampal pairs in controls and in 36% of possible pairs in MAM (p=0.002). There were no significant group differences in PFC to PFC pairs with edges identified in 46% of all possible pairs in controls and 41% of all possible pairs in MAM. Remarkably, there were significant CPFs for pairs spanning the hippocampus and PFC, indicating tightly coordinated firing across structures, including feedback communication from the PFC to the hippocampus. Hippocampus to PFC pairs had edges in 64% of possible control and 60% of possible MAM pairs. PFC to hippocampus pairs had edges in 67% of possible control and 58% of possible MAM pairs. There were no significant group differences.

Pairs of cells were then categorized into those in which both cells in the pair encoded the sample phase (*sample-encoding pairs*) and those pairs that did not. Similarly, pairs were also categorized into those in which both cells significantly differentiated correct from incorrect lever presses at the match phase (*match-encoding pairs*) and those pairs that did not. Both sample- and match-encoding pairs had a significantly higher proportion of edges between them compared to pairs of cells in which at least one cell did not significantly encode either sample or match (Table 1).

**Table 1.**
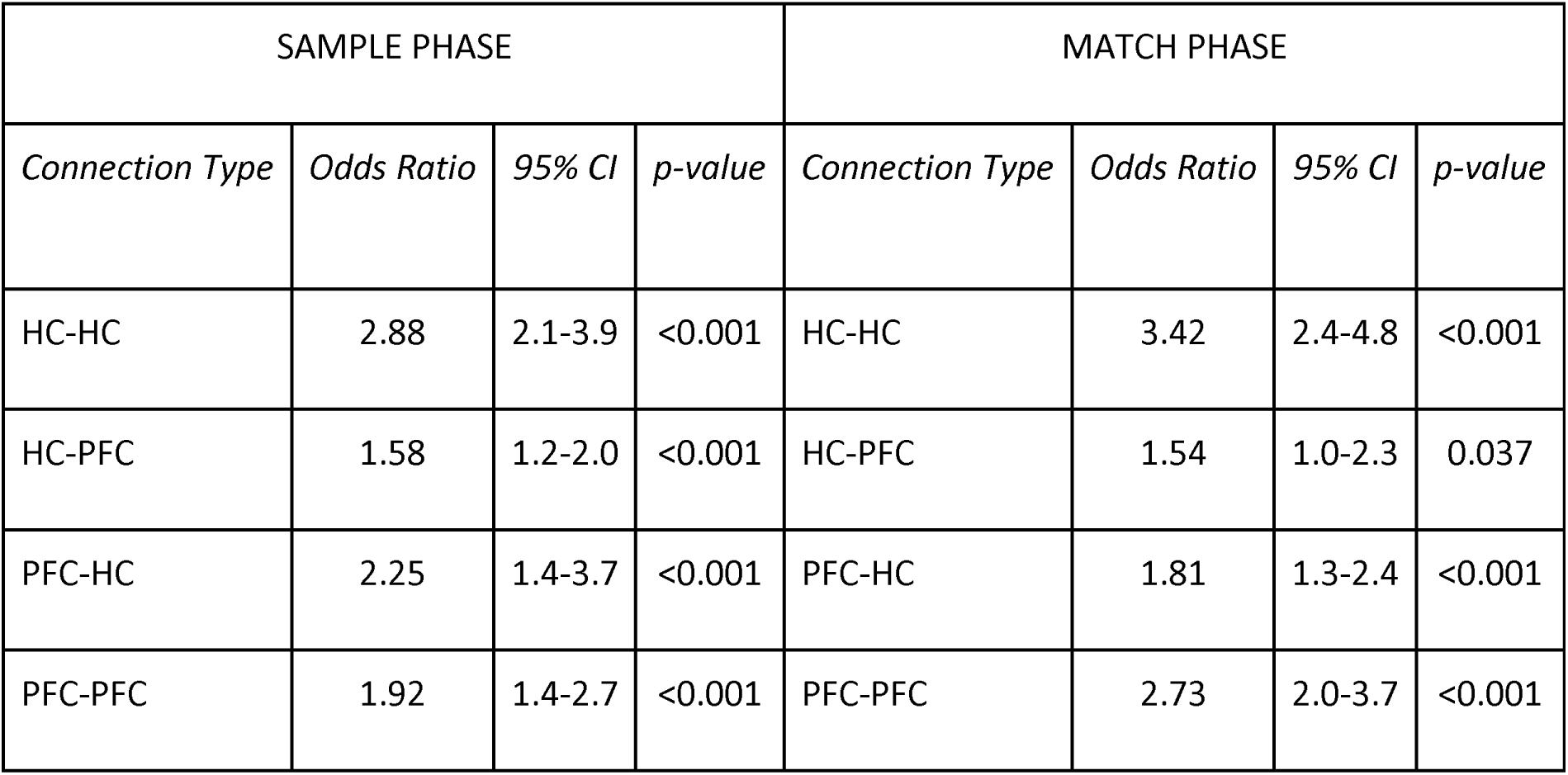
Neurons within and between brain regions encoding task parameters are more likely to be connected than neurons not encoding either the sample or the match phase (HC: hippocampal neuron, PFC: prefrontal cortical neuron).

| SAMPLE PHASE |  |  |  | MATCH PHASE |  |  |  |
| --- | --- | --- | --- | --- | --- | --- | --- |
| *Connection Type* | *Odds Ratio* | *95% CI* | *p-value* | *Connection Type* | *Odds Ratio* | *95% CI* | *p-value* |
| HC-HC | 2.88 | 2.1-3.9 | <0.001 | HC-HC | 3.42 | 2.4-4.8 | <0.001 |
| HC-PFC | 1.58 | 1.2-2.0 | <0.001 | HC-PFC | 1.54 | 1.0-2.3 | 0.037 |
| PFC-HC | 2.25 | 1.4-3.7 | <0.001 | PFC-HC | 1.81 | 1.3-2.4 | <0.001 |
| PFC-PFC | 1.92 | 1.4-2.7 | <0.001 | PFC-PFC | 2.73 | 2.0-3.7 | <0.001 |

To further explore the relationship between functional network structure and task encoding, we analyzed the node degree (i.e. the number of connections a cell has) of cells in the functional network. To compare across differently sized networks, we rank-normalized degrees. Because edges in the network are directional, each node has an *in-degree* and *out-degree*, corresponding to the number of incoming and outgoing connections. In all cases, the cells that significantly coded at the sample or match had higher in- and out-degree (p=0.001 for both in-degree and out-degree; Fig. 7a,b), demonstrating that the more integrated a cell is into the functional network, the more likely it is to encode the SWM task parameters. Importantly, the degree measures did not distinguish whether the cells in the ensemble were from hippocampus or PFC. Thus, the hippocampal-PFC ensemble in its entirety is important for performance (Fig. 7a,b). These data demonstrate that the functional organization of ensembles of cells at short-timescale predicts the generation of long-timescale task-related dynamics and SWM task performance.

**Figure 7.**
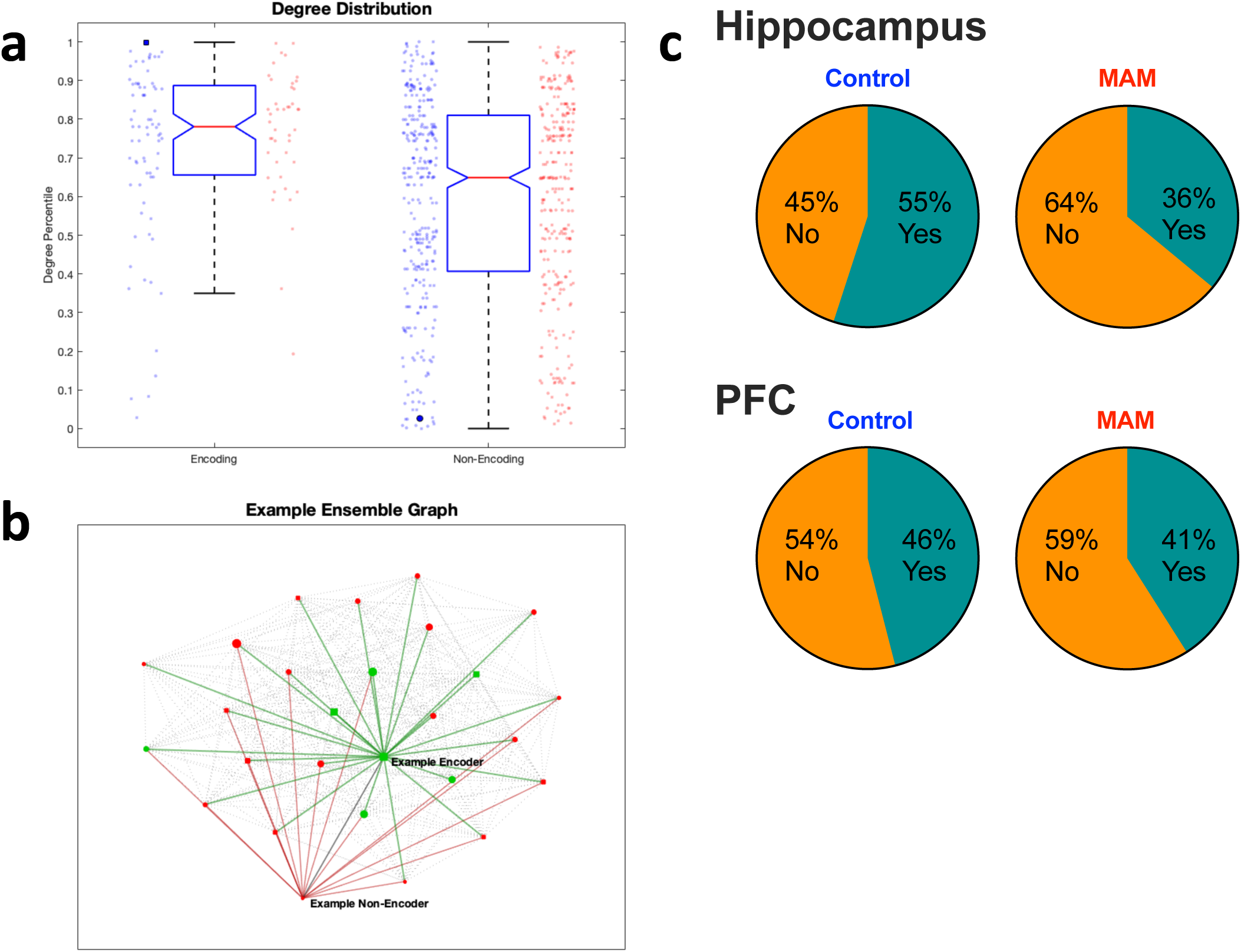
Neurons that encode task parameters are more densely functionally connected in both groups. Median degree for each neuron that is encoding task parameters vs non-encoding neurons, with individual values of neurons from controls (blue) and MAM animals (red) shown on either side of the box plot of both group averages (a), 25% and 75% percentile shown in the box with whiskers extending to the most extreme data points. Plot of the ensemble from which the darkened data points in (a) are acquired is shown in (b), with encoding neurons labeled in green and non-encoders labeled in red. Networks are densely connected but not all neuron pairs are functionally connected. Size of each node represents the total number of connections and shows that, on average, the encoders are better connected. Square nodes are neurons are from hippocampus and circular nodes are neurons from PFC in (b), showing the functional interconnectedness of both structures. On average, fewer pairs of neurons in the hippocampus (top, c) and the PFC (bottom, c) are significantly connected in MAM animals compared to controls.

To establish whether timing and population dynamics were independent predictors of task performance, we modeled performance with a multivariable regression as a function of group, mean task length, mean PC1 scores from both hippocampus and PFC, proportion of edges, proportion of neurons that encode the sample and match phase and their interactions. This final model requires all of the main effects and some of the interactions, and accounts for 79% of the variance in performance, compared to 54% by a model using rate dynamics alone (p=0.037; Fig. 8a). Therefore, the firing dynamics of neurons encoding the sample lever position and leading up to the match phase decision point *<u>and</u>* oscillatory timing and functional network structure independently predict overall performance, as the full model accounts for more of the variance than can be explained by rate encodings alone.

**Figure 8.**
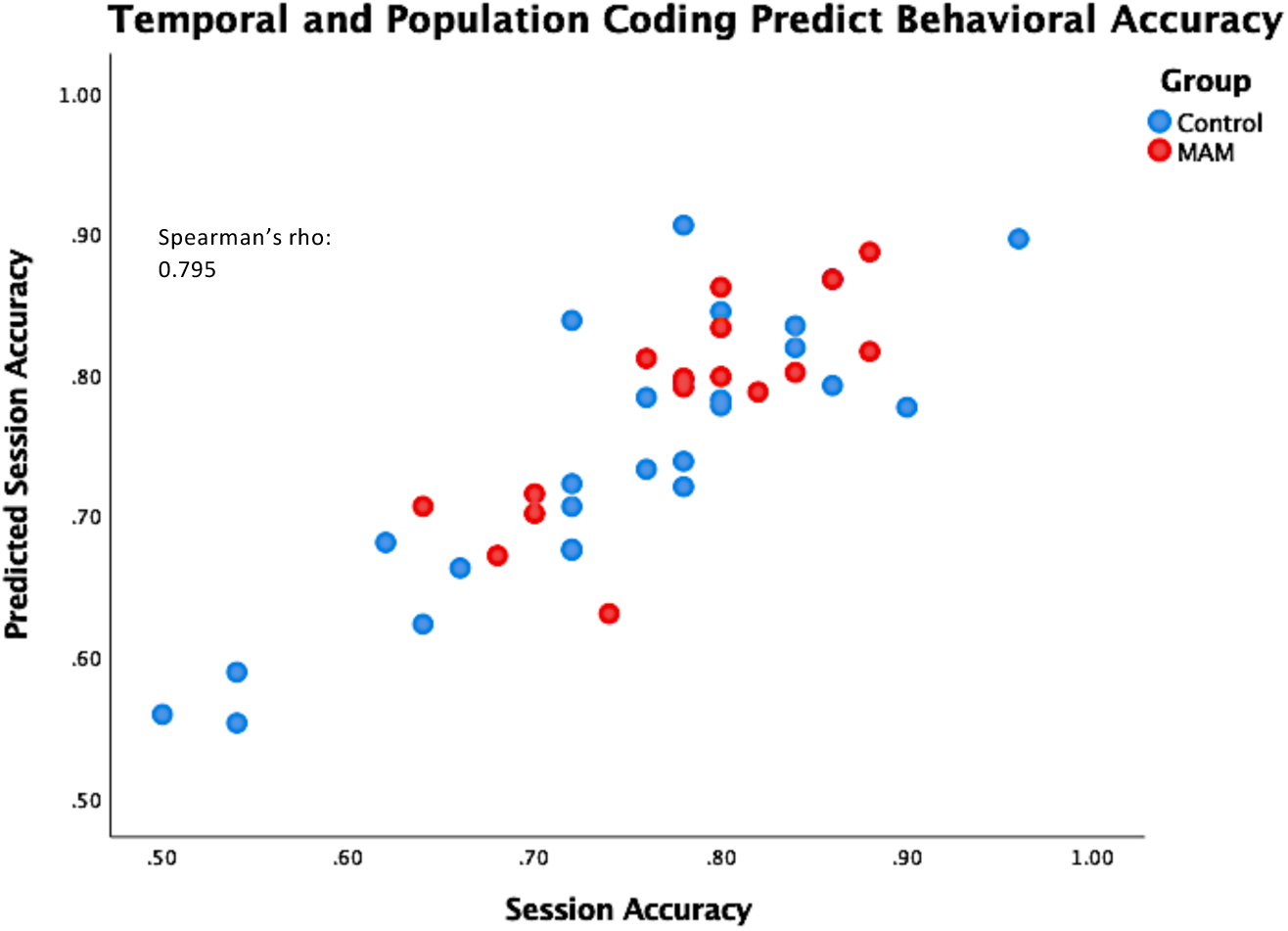
Neuronal dynamics in PFC and hippocampus predict behavioral accuracy in the SWM task in MAM rats and controls. Overall session accuracy (number of correct trials/number of incorrect trials) was predicted using a GEE that contained only rate, temporal and population coding parameters. Predicted session accuracy from these coding parameters was very well correlated with actual session accuracy, indicating that these coding parameters of neurons in the PFC and the hippocampus account for roughly 80% of the variance in measured accuracy.

## Discussion

The major finding in this study is that timing dynamics of ensembles of neurons spanning the hippocampus and PFC are directly related to the intermittent rate encoding dynamics of SWM. The hippocampal neurons with strong oscillatory modulation and the hippocampal and PFC neurons that are tightly integrated into the functional network are the same neurons that encode the sample lever position and are involved in the choice at the match phase of a DNMS task. In the context of a developmental brain disorder, there are fewer neurons that encode the task, and these neurons have lower oscillatory fidelity and are less tightly integrated into the functional network. Importantly, the oscillatory and functional network parameters, both of which capture ongoing dynamics at sub-second timescales, are independent predictors of behavioral performance. These results show that the ability of a neural circuit to support emergent, coordinated spiking dynamics throughout the hippocampal-prefrontal network is a system-level mechanism explaining performance in a SWM task and explaining deficits in animals with malformed neural circuits.

In the DNMS task, the hippocampus encodes the lever position (left vs. right) and transmits this information to the PFC where it is stored for decision-making^5^. The PFC, in return, receives this information and integrates it with internal states representing, for example, attention, motivation, the organism’s high-level goals (correct vs. incorrect) and possible motor strategies (press left vs press right) for achieving those goals^32^. However, there are also large numbers of hippocampal neurons with firing that significantly discriminates correct from incorrect presses at the match, and PFC neurons whose firing discriminates left from right lever position at the sample (Fig. 3e,f). This finding agrees with a model in which information is multiplexed across these structures. Our data demonstrate that, even though the cognitive roles of hippocampus and PFC are distinct, they each process all task parameters, both at the level of cognitive rate coding and its relationship to task performance, in normal rats and those with a brain malformation. However, only a subset of neurons is rate coding at decision points, raising the question of what determines which cells are in the coding subset. Because of the established dynamic interaction between the hippocampus and PFC at the level of the LFP throughout an SWM trial, we hypothesized that oscillatory fine spike timing dynamics and functional connectivity between structures were important determinants of cognitive encodings.

The encoding and transfer of information during rhythmic oscillations in the brain has been repeatedly implicated as a critical mechanism for multiple forms of learning and memory^33^. In particular, in the hippocampus, the microcircuits of CA1 pyramidal cells oscillate at approximately 8 Hz^29^, and the fidelity of a cell’s firing within population-level theta oscillations is known to be important for the encoding of spatial memory^17^. We extend this observation to SWM, as strongly theta-modulated neurons are precisely those that significantly encode the SWM task parameters. At the population level, our data demonstrate that fine spike timing between neurons in the hippocampus and PFC predicts pairs of cells in which both are encoding at the sample and match phases, i.e. multiplexing of task parameters (Fig. 7). This effect is highly non-trivial, as a functional network edge between a pair of cells is defined over-and-above the tendency of cells to either have oscillatory auto-correlation or to co-modulate with other cells in the ensemble. Thus, multiplexing of task parameters during working memory is directly related to the fidelity of timing characteristics between specific pairs of cells, including pairs that span structures. Importantly, the functional networks display feedback from the PFC to the hippocampus. In contrast to a feed-forward model from hippocampus to PFC, in which the hippocampus encodes spatial information to be subsequently processed by the PFC, we observe ongoing dynamical coordination of hippocampal cells subsequent to PFC firing. Because PFC neurons do not synapse directly onto CA1 neurons, other structures, e.g. the nucleus reuniens, which has been implicated as a relay during SWM^34–36^, are likely required to coordinate co-firing across the hippocampal-prefrontal network^37^. The relationship between such relay structures and the CPFs in our functional networks is an important avenue for future studies. Taken together, these results implicate a model in which ongoing oscillatory coordination among neurons in the hippocampal-prefrontal network defines a functional network that is poised to receive sensory inputs in the hippocampus and process them as working memory through dynamical coordination throughout the hippocampal-prefrontal network.

Translational systems neuroscience attempts to identify the properties of neural networks that impair the ability of the brain to carry out cognitive computations with the ultimate goal of identifying therapies based upon manipulating those properties. To this end, we have shown that in the MAM model, a clinically-relevant model of structural disorganization of the hippocampal and PFC circuits, alterations in oscillatory and functional network dynamics directly predict cognition and SWM task performance. Given that overall oscillatory disruptions are observed in diseases from schizophrenia to Alzheimer’s disease and epilepsy, our results may have important generalizability across a variety of brain diseases associated with working memory impairments. We demonstrate that, on a per cell basis, cognitive encoding can be directly predicted from oscillatory behavior within an ensemble, and that this relationship accounts for the cognitive deficits in MAM animals. MAM networks, on average, had fewer encoding cells, which were less precisely temporally modulated, suggesting that although the MAM neurons can accurately subserve working memory, their networks are doing so less efficiently. We suspect this low efficiency computation may be responsible for the significantly longer time to complete trials in MAM animals. Despite these differences, however, our data show that the relationship between rate, oscillatory, and functional network dynamics is maintained in a structurally malformed brain. This provides a potential system-level mechanism for the recent clinical observation in patients with epilepsy showing that a non-specific 10-200 Hz electrical stimulation for 500 ms during a working memory task improves working memory performance, specifically when the subject’s brain was predicted to be in a “non-encoding” state^38^. In that case, perturbing neural dynamics with a short wavelength oscillation was sufficient to engage the brain into a receptive state for correctly perceiving and retaining information in working memory. However, these stimulations were time-locked to the task, and may not represent a viable therapeutic option, as there is no obvious trigger for stimulation outside of a controlled experiment. Whether it is possible to use background brain stimulation to normalize oscillatory and functional network dynamics in structurally disorganized neural circuits is still an open question. However, our results suggest that this may be a viable strategy to normalize working memory, as the oscillatory and functional network dynamics were measured throughout the experimental session, and therefore represent a scaffold on which intermittent working memory stimuli are incorporated as memories.

## Methods

### Contact for Reagent and Resource Sharing

Further information and requests for resources and reagents should be directed to and will be fulfilled by the Lead Contact, Matt Mahoney.

### Experimental Model and Subject Details

#### Animals

All animal procedures were approved by the University of Vermont Institutional Animal Care and Use Committee, under United States Department of Agricultural and Association for the Assessment and Accreditation of Laboratory Animal Care International approved conditions, in accordance with National Institutes of Health guidelines. Time-mated pregnant Sprague-Dawley rat dams were randomly selected for intraperitoneal injection with either 20 mg/kg MAM (Midwest Research Institute Global, Kansas City, MO) or saline at embryonic day (E) 17. Male and female offspring were housed with a 12 h light/dark cycle and ad libitum access to food and water until behavioral studies were initiated. Studies began when animals were p45. All animals were group housed until experiments commenced to allow for food deprivation protocol three days prior to initiation of behavioral experiments. Animals were food deprived to approximately 85% of starting weight.

### Method Details

#### Surgical implantation of electrodes

Rats were anesthetized with isoflurane (2-3% in oxygen) and placed in a stereotaxic frame (Kopf Instruments, Tujunga, CA). Custom-built electrodes containing four independently drivable tetrodes were implanted 3.2 mm posterior to the bregma, 2.2 mm lateral and 1.7 mm deep into the dorsal CA1 hippocampus and 1.5 mm anterior to bregma, 0.5 mm lateral and 2.5 mm in control animals. Coordinates were adjusted by 10% for MAM animals to account for smaller brain size as has been previously published^17,39^. Tetrode location was visually verified in PFC and hippocampus post-mortem in all animals.

#### Data Acquisition and post-processing

Tetrode assemblies were advanced 50 µm twice a day until hippocampal theta oscillations (6-12 Hz), sharp waves and ripples were observed in the EEG. Electrodes were then advanced in 25 µm increments until CA1 single unit activity was detectable. Single unit activity was recorded when waveforms above 40 µV in amplitude were observed on one or more tetrodes. The signal from the electrodes was preamplified directly from the rat’s head by operational amplifiers and transmitted via a custom cable to a Neuralynx recording system (Neuralynx, Bozeman, MT). Signals were recorded at 33.3kHz sampling frequency wide-band and then subsequently filtered from 500-9000Hz, thresholded at at least 3x RMS noise in the first 20 s of the signal. Putative single-unit firing was identified by clustering action potentials from this filtered and thresholded signal using Neuralynx Spike Sort 3D (Bozeman, MT).

#### Behavioral Task

An operant box (Med Associates Inc., St. Albans, VT) enclosed in a sound-attenuating chamber was used for delayed non-match to sample (DNMS) experiments. Inside the box, two retractable levers were located on one wall separated by a pellet dispenser. The opposite wall contained a nose-poke hole. Stimulus lights were located above each lever, the food cup and the nose-poke hole. At p45, rats were food deprived to approximately 85-90% of starting weight and were rewarded during the task with sucrose pellets from the pellet-feeder. Behavioral software (Med Associates Inc., St. Albans, VT) reported the number of correct trials completed in the entire session. For all behavioral experiments, animals were acclimated to the testing room for at least 2 hours before testing. The task involved several training steps before a full trial could be executed. Once criterion was achieved in these initial phases, the DNMS sessions began. DNMS trials involved the presentation of a sample lever, which the animal had to press, followed by a nose-poke, followed by the “non-match phase”, which involved the presentation of two levers from which the animal had to select the lever that was not pressed during the sample phase (Figure 1). DNMS was done in blocks with increasing delay times between the nosepoke and non-match phases; block 0 has a 0 second delay, block 1 has trials with 0-5 second delays, block 2 has trials with 0-10 second delays, and block 2A has trials with 5-10 second delays. The inter-trial interval was set at 10 seconds. Criteria to pass to the next block include accuracy of 80% or more (40 or more correct trials) during the session and no clear lever preference. Lever preference or less than 60% accuracy (fewer than 30 correct trials) resulted in regression to the previous block or phase of DNMS training. Behavioral software (Med Associates Inc., St. Albans, VT) reported number of presses on each lever, number of rewards dispensed, and number of nose-pokes. Food rewards were used in all tasks (45mg Noyes food pellet; Research Diets Incorporated, New Brunswick, NJ).

#### GLMs - Event filters

We binned rasters of neural spike trains at *Δ* = 1 ms to obtain binary vectors (*t*) whose value is 1 if a spike occurred at time *t* and 0 otherwise. We modeled *r*(*t*) as a Poisson random variable with a time varying mean firing rate *λ*(*t*). The log-likelihood function for this model is

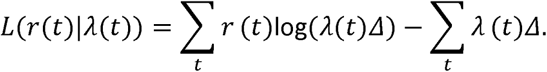

To account for rate modulation around lever presses, we modeled *λ*(*t*) as

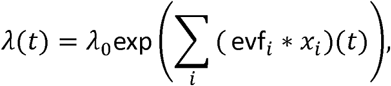

where *λ*_0_ is the baseline firing rate, evf_*i*_(*t*) is an *event filter* function, *denotes convolution, and *x*_*i*_(*t*) is a raster of external events (e.g., left, right, or sample lever presses at the sample phase; *cf.*). The function evf_*i*_ encodes the firing rate modulation around the event *x*_*i*_. For the sample phase, evf_*i*_ is non-zero only for *t* > 0, encoding post-lever-press modulation, while at the match phase evf_*i*_ is non-zero only for *t* < 0, encoding pre-lever-press modulation. We parametrized the filters evf_*i*_ (*t*) using 5 *raised cosine* basis functions using Matlab scripts from Pillow *et al.* ^26^ (ihprs.ncol = 5; ihprs.peaks = [0.05 2]; ihprs.b = 0.5; Fig. S3 These functions describe the rate modulations over the ∼2 s after a lever press. With this basis each filter, evf_*i*_, is represented as evf_*i*_(*t*) = Σ_*k*_ *β*_*k*_ rc_*k*_(*t*). With this parametrization, the log-likelihood becomes a function of the *β*_*k*_’s,*L*(*r*_t_|*β*_*k*_). We estimated the coefficients *β*_*k*_ using maximum likelihood

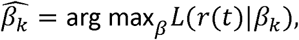

using the Matlab function `fminunc`. The value

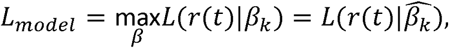

is the *model log-likelihood.*

At the sample phase, we generated two models: a *left-right model* with distinct filters for left and right lever presses, and a *sample null* model with a single filter for both right and left lever presses. We did the same at the match phase to make a *correct-incorrect model* and a *match null model*.

#### GLMs – post-spike filters

To account for post-spike rate modulation (auto-correlation), we modeled *λ*(*t*) as

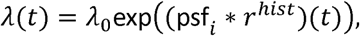

where *r*^*hist*^ denotes the spiking history of the cell and psf_*i*_ is a *post-spike filter*, which encodes the firing rate modulation after a spike. We parametrized the filters psf_*i*_(*t*) using 10 raised cosine basis functions (ihprs.ncol = 10; ihprs.peaks = [0.01 0.5]; ihprs.b = 0.5; Fig. S3 and one immediate post-spike impulse to capture the refractory period (*cf.* ^17^) To avoid overfitting the PSF’s, we added a *ridge penalty* to the log-likelihood function

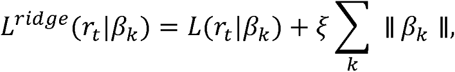

where *ξ* is a hyper-parameter determines the strength of the ridge penalty. We selected *ξ* using evidence maximization, as described in Park *et al*.^*27*^

#### GLMs - coupling filters

To account for functional interactions between cells (cross-correlation), we modeled *λ*(*t*) as

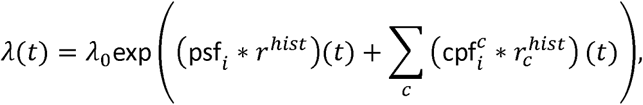

where psf_*i*_ is as above, 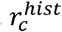 denotes the spiking history of the cell *c* and 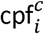 is a *coupling filter*, which encodes the firing rate modulation after a spike. We parametrized the filters 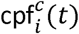 using 6 raised cosine basis functions (ihprs.ncol = 6; ihprs.peaks = [0.001 0.5]; ihprs.b = 0.5; Fig.S3. We fit these *ensemble models* identically to the PSF-only models.

#### Log-likelihood ratios tests, task-parameter encoding, and functional networks in ensembles

To establish whether a neuron distinguished between left vs. right at the sample, we computed the difference of the log-likelihoods for the fitted left-right and sample null models to obtain *log likelihood ratios* (LLRs)

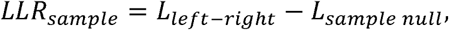

which is equivalent to taking the logarithm of the ratio of model likelihoods. We define *LLR*_*match*_ similarly. Under the null hypothesis, the LLR’s are *χ*^2^-distributed with 5 degrees of freedom, corresponding to the difference in the number of parameters in the models. A neuron was classified as encoding left vs. right or correct vs. incorrect if the corresponding LLR had a *p*-value less that 0.05 after false discovery rate correction for multiple hypotheses.

Similarly, to define functional networks among neurons, each ensemble model was fit with and without each coupling filter and the corresponding LLR was computed. Under the null hypothesis, these LLR’s are *χ*^2^-distributed with 6 degrees of freedom, corresponding to the number of parameters defining the coupling filter. A coupling filter was considered significant if its *p*-value (after false discovery rate correction) was less than 0.05. Graphically, we can visualize a significant coupling filter as a *directed edge* in a functional network (*cf* Fig. 7b).

#### Match filter differences

To determine the differences in the firing regulation before correct vs. incorrect lever presses at the match, we took the difference between the correct and incorrect event filters

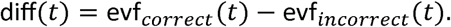

We normalized each filter to have sum-of-squares equal to one prior to taking the difference. We then performed an uncentered principal components analysis (i.e. a singular value decomposition; Matlab command ‘svd’) of the filter differences separately by region and group to identify the dominant patterns among the filter differences. In order to visualize these differences across cells in a heat map, we sorted the filter differences by their projection onto the first principal component.

### Quantification and Statistical Analysis

All analyses were carried out using SPSS v25 within a generalized estimating equation (GEE) framework. Means (or odds ratios), confidence intervals and p-values are reported in the results section of the text. GEE allows us to account for multiple observations from single animals, assume the correct distribution for the data and apply the correct link function. Ns were the number of animals per group (N=6 control, N=7 MAM rats), adjusted for multiple cells over the course of multiple sessions per animal. Transformation to a normal distribution was performed when possible. An exchangeable working correlation matrix was used as a default. Goodness of fit was evaluated using the Corrected Quasi-likelihood under Independence Model Criterion (QICC) with the lowest value model being used as final. For normally distributed data we also visually assessed residuals plotted against predicted values and plotted residuals on a Q-Q plot to confirm normality.

For the evaluation of the working memory aspects of the DNMS trial we compared the proportion of correct trials as a function of all completed trials across groups. The time from sample lever press to match lever press was calculated for each completed trial and compared between groups, and between correct and incorrect trials.

Log likelihood ratios were calculated from event filters with dynamic changes in the range of seconds as above. The data were binarized to define neurons that significantly distinguished right from left lever at the sample phase and correct from incorrect presses at the match phase, using a logistic regression approach. The proportion of significantly coding cells were compared across groups in each brain region independently. We then defined cells by whether they significantly coded both the sample and match phases and compared these proportions between groups in each brain region. Finally, we used behavioral performance as the dependent variable with proportion of cells that significantly code both sample and match phases and trial length as independent variables to determine whether coding characteristics were related to behavioral outcomes.

Short timescale (millisecond) dynamics were evaluated using post-spike filters as described above. PC 1 scores of the PSF were converted to *normal scores* by taking the inverse normal cumulative distribution function of their normalized ranks^31^. The first principal component (PC1) score, was used as the dependent variable and compared across groups in each brain region independently. To establish the relationship between cells that significantly encode in the second timescale (rate encoding) and overall millisecond firing dynamics (timing coding) we used the binarized LLR data as the dependent variable and the first principal component normal score as the independent variable and report the odds ratios as a function of standard deviation change in PC1 normal scores.

To characterize population dynamics we generated coupling filters between all cells across the entire session in the millisecond timescale. The data were binarized into couples that show significant comodulation (defined using Granger causality) and those that do not. The proportion of edges as a function of the total number of possible edges was used as the dependent variable and compared across groups. Hippocampus to hippocampus, prefrontal cortex to prefrontal cortex, hippocampal to prefrontal cortex and prefrontal cortex to hippocampus filters were evaluated independently. To establish the relationship between coding characteristics and comodulation we binarized pairs of cells into those in which both cells significantly coded the sample phase and those in which at least one cell did not have a significant log-likelihood ratio. We repeated this at the match phase. These data were the dependent variable and we included the proportion of edges as the independent variable for sample and match phases independently. Normalized node degree was evaluated as a network parameter as described above. As this is directional we evaluated the normalized in-node degree and normalized out-node degree as dependent variables separately. The node degree was the dependent variable with significant log-likelihood ratios from event filters as the independent variable.

The final analysis modeled performance with a multivariable regression as a function of group, mean task length, mean PC1 scores from both hippocampus and PFC, proportion of edges, proportion of neurons that encode the sample and match phase and their interactions. behavioral performance as the dependent variable with proportion of cells that significantly code both sample and match phases and trial length as independent variables. The correlation coefficient obtained from this analysis was compared to the analysis with only task dependent parameters as independent variables using a Fisher transform and then a Z-test.

---

**Supplemental Figure 1.**
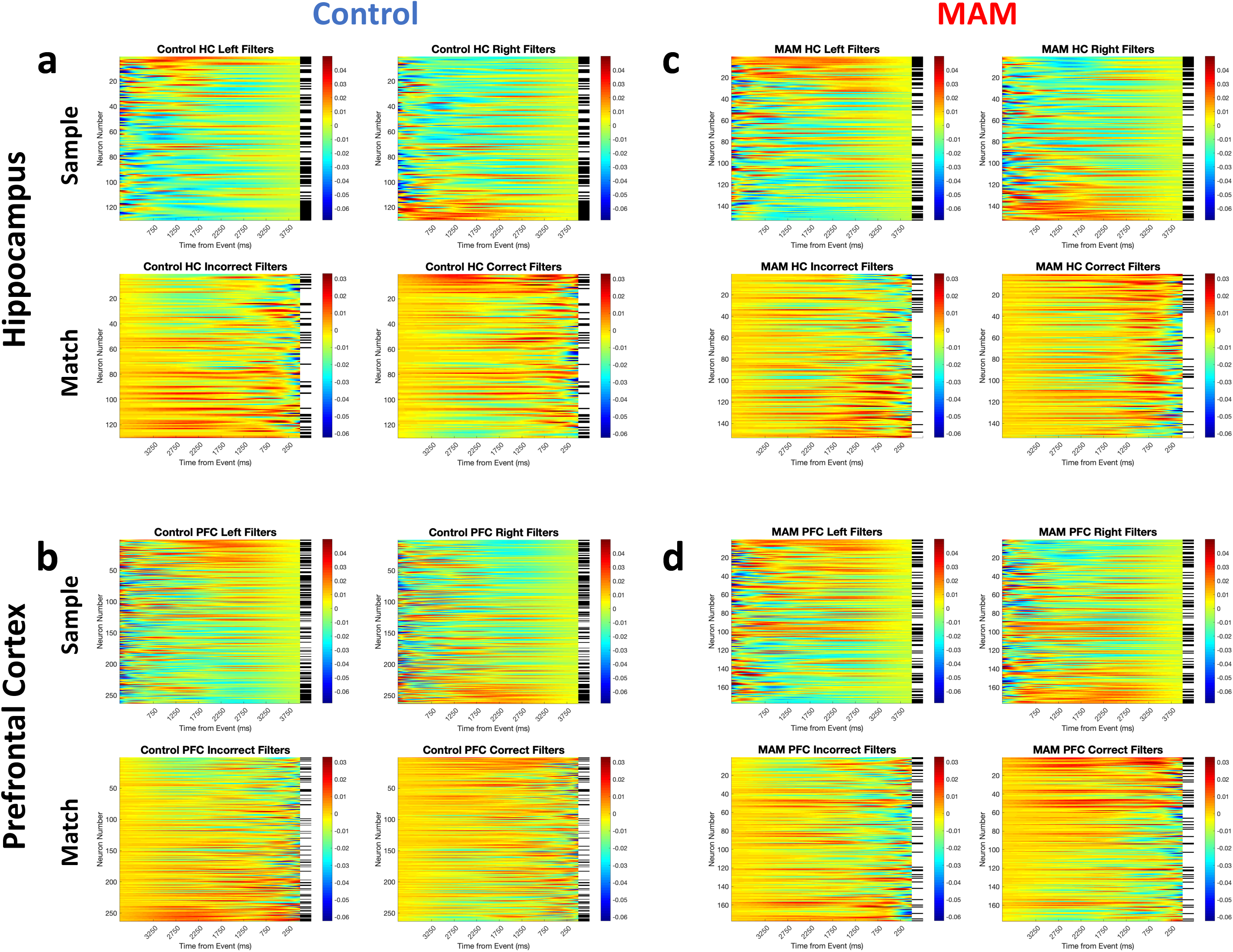
Event-related firing is heterogeneous. Heatmaps show event filters of neuronal firing either 4s after the sample press (top panels) or 4 s before the match press (bottom panels) in hippocampus (**a**) or PFC (**b**) in controls, and hippocampus (**c**) or PFC (**d**) of MAM animals. Significant firing heterogeneity around the task parameters, including persistently firing neurons, can be seen in accordance with previously published work.

**Supplemental Figure 2.**
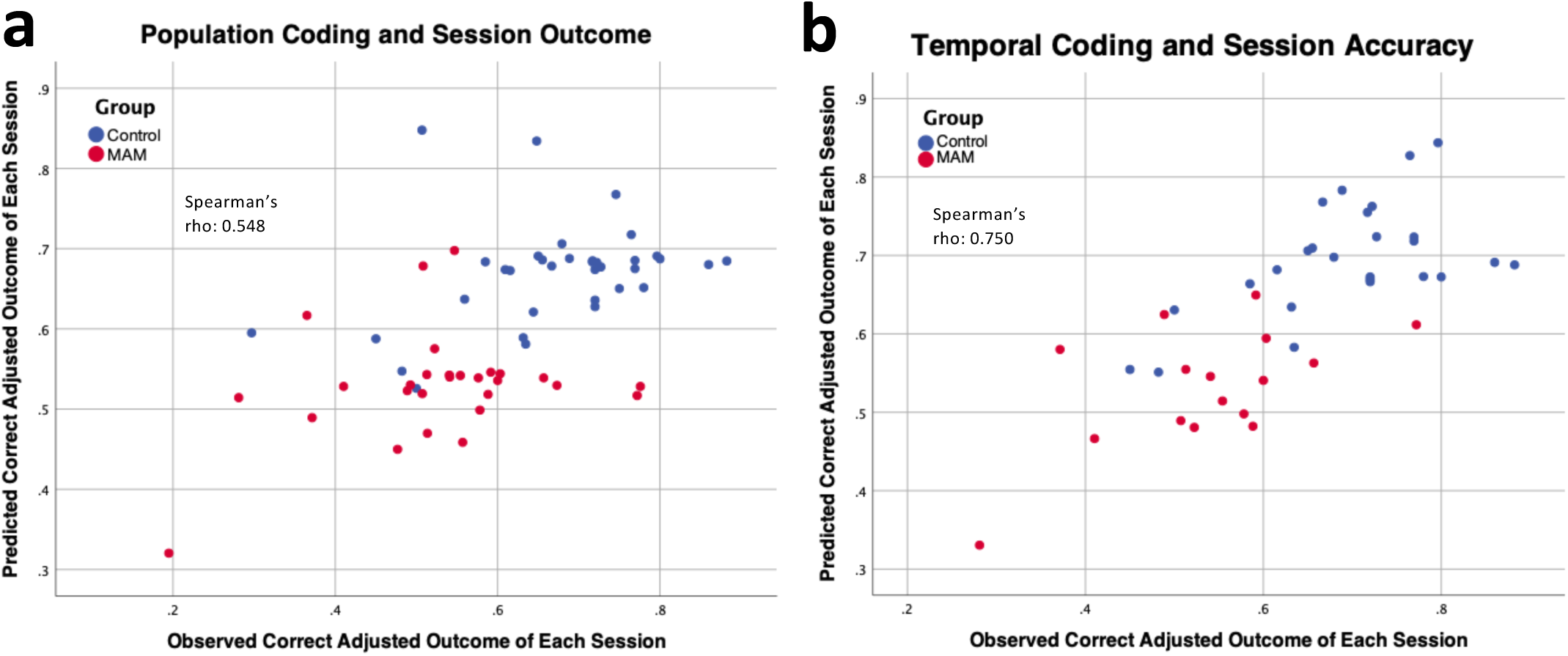
Population and temporal coding parameters also predict session accuracy. A model that predicts session accuracy from only event coding parameters, i.e. the proportion of neurons in a recording session that are encoding the task parameters, (**a**) significantly correlates with observed accuracy in the task. A model that predicts session accuracy from only temporal coding parameters (**b**) also significantly correlates with observed session accuracy in the task.

**Supplemental Figure 3.**
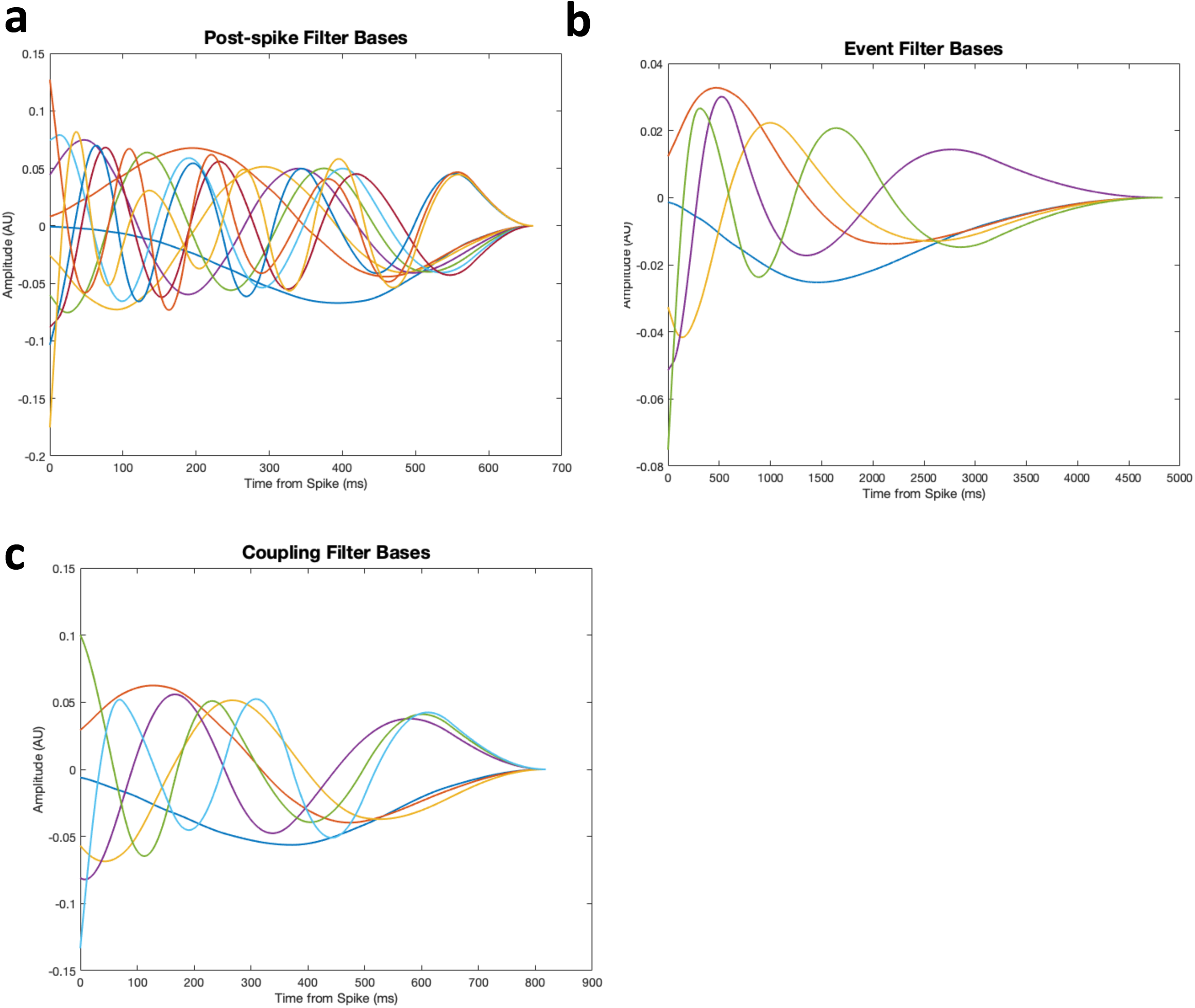
Basis functions for GLM. Basis functions for post-spike filters (**a**), event filters (**b**) and coupling filters (**c**).

